# DLL4-Notch3-WNT5B axis is a novel mediator of bi-directional pro-metastatic crosstalk between melanoma and lymphatic endothelial cells

**DOI:** 10.1101/2023.04.21.537773

**Authors:** Sanni Alve, Silvia Gramolelli, Joonas Jukonen, Susanna Juteau, Anne Pink, Atte Manninen, Elisa Monto, Madeleine H. Lackman, Olli Carpén, Sinem Karaman, Pipsa Saharinen, Kari Vaahtomeri, Päivi M. Ojala

## Abstract

Despite strong indications that melanoma interaction with lymphatic vessels actively promotes melanoma progression, the molecular mechanisms are not yet completely understood. To characterize molecular factors of this crosstalk we established human primary lymphatic endothelial cell (LEC) co-cultures with human melanoma cell lines. Here, we show that co-culture with melanoma cells induced transcriptomic changes in LECs and led to multiple alterations in their function. WNT5B, a paracrine signaling molecule upregulated in melanoma cells upon LEC interaction, was found contributing to the functional changes in LECs. Moreover, *WNT5B* transcription was regulated by Notch3 in melanoma cells following the co-culture with LECs, and Notch3 and WNT5B were co-expressed in melanoma patient primary tumor and metastasis samples. Moreover, melanoma cells derived from LEC co-culture escaped efficiently from the primary site to the proximal tumor draining lymph nodes, which was impaired upon WNT5B depletion. This supports the role of WNT5B in promoting the metastatic potential of melanoma cells through its effects on LECs. Finally, DLL4, a Notch ligand expressed in LECs, was identified as an upstream inducer of the Notch3-WNT5B axis in melanoma. This study elucidates WNT5B as a novel molecular factor mediating bi-directional crosstalk between melanoma cells and lymphatic endothelium and promoting melanoma metastasis.

## INTRODUCTION

Metastatic melanoma is the most lethal form of skin cancer, and its occurrence continues to raise especially in the western population. The prognosis for patients with metastasized disease is poor with the long-term survival rate of only 10% (1). It is estimated that 80% of the melanoma metastases spread from the primary tumor to distant sites through the lymphatic vasculature. The importance of lymphatic vasculature for melanoma metastasis is further supported by observations that the peritumoral lymphatic vessel infiltration correlates with higher metastatic rate and thereby increased mortality in melanoma (2).

Tumors actively shape and modify the surrounding lymphatic system. Using 3D imaging for a mouse pancreatic cancer model, it was shown that the developing tumor caused extensive changes in the architecture of tumor-associated lymphatic vasculature, including remodeling of the existing lymphatic vasculature, invagination of the endothelium and vasodilation (3). For melanoma it has been demonstrated that cells can release lymphangiogenic factors to locally control lymphatic vasculature (4). Tumors not only cause changes in the local lymphatic vasculature but also in the tumor draining lymph nodes; these lymph nodes appear larger due to proliferation of the resident macrophages and stromal lymphatic endothelial cells (5). Furthermore, melanoma-secreted extracellular vesicles (EVs) educate the tumor draining lymph nodes to enhance lymphatic metastasis in many ways, for instance through neural growth factor (NGF) receptor-dependent signaling (6), by shuttling tumor antigens to lymph node LECs for cross-presentation on MHC-I resulting in apoptosis induction in antigen-specific CD8^+^ T cells (7), or by compromising maturation process of the lymph node residing dendritic cells (8).

Not only is melanoma actively remodeling lymphatic vessels to promote cancer progression, but there is also increasing evidence that lymphatic vasculature can in turn directly modify the properties of melanoma cells, which suggests that active, bidirectional crosstalk occurs between the cancer cells and the lymphatic vasculature. For example, tumor associated lymphatic endothelial cells (LECs) can actively attract the cancer cells, leading to enhanced cancer cell migration towards the lymph vessels (9, 10). In addition, we have previously shown that direct contact of melanoma cells with LECs strongly augments melanoma invasion and metastasis through a matrix-metalloproteinase-14 (MMP14, also known as MT1-MMP) dependent induction of Notch3 in the melanoma cells (11).

While the pivotal role of the lymphatic vasculature in promoting cancer dissemination is well conceded, the initial steps of lymphogenic cancer metastasis and in particular the nature and molecular signaling involved in tumor cell communication with the surrounding lymphatic vasculature, are not yet completely understood. To better understand this reciprocal, metastasis-promoting crosstalk we implemented co-culture systems of melanoma and LECs to discover molecular determinants for the communication between these two cell types. We found that melanoma cells induced metastasis-promoting, molecular and functional changes in LECs mediated by a novel DLL4-Notch3-WNT5B signaling axis.

## RESULTS

### Melanoma induces functional and transcriptional changes in LECs

To characterize possible changes in LEC upon their co-culture with melanoma cells, we used a combination of 2D and 3D functional assays to assess the LEC properties. For each of these experiments, LECs were cultured either in monotypic control cultures or co-cultured with a panel of GFP expressing melanoma cell lines: both metastatic (WM852 and WM165) and non-metastatic (WM793). After two days in co-culture, cells were sorted and the LECs used for subsequent assays (indicated as LEC*; Figure 1A).

**Figure 1.**
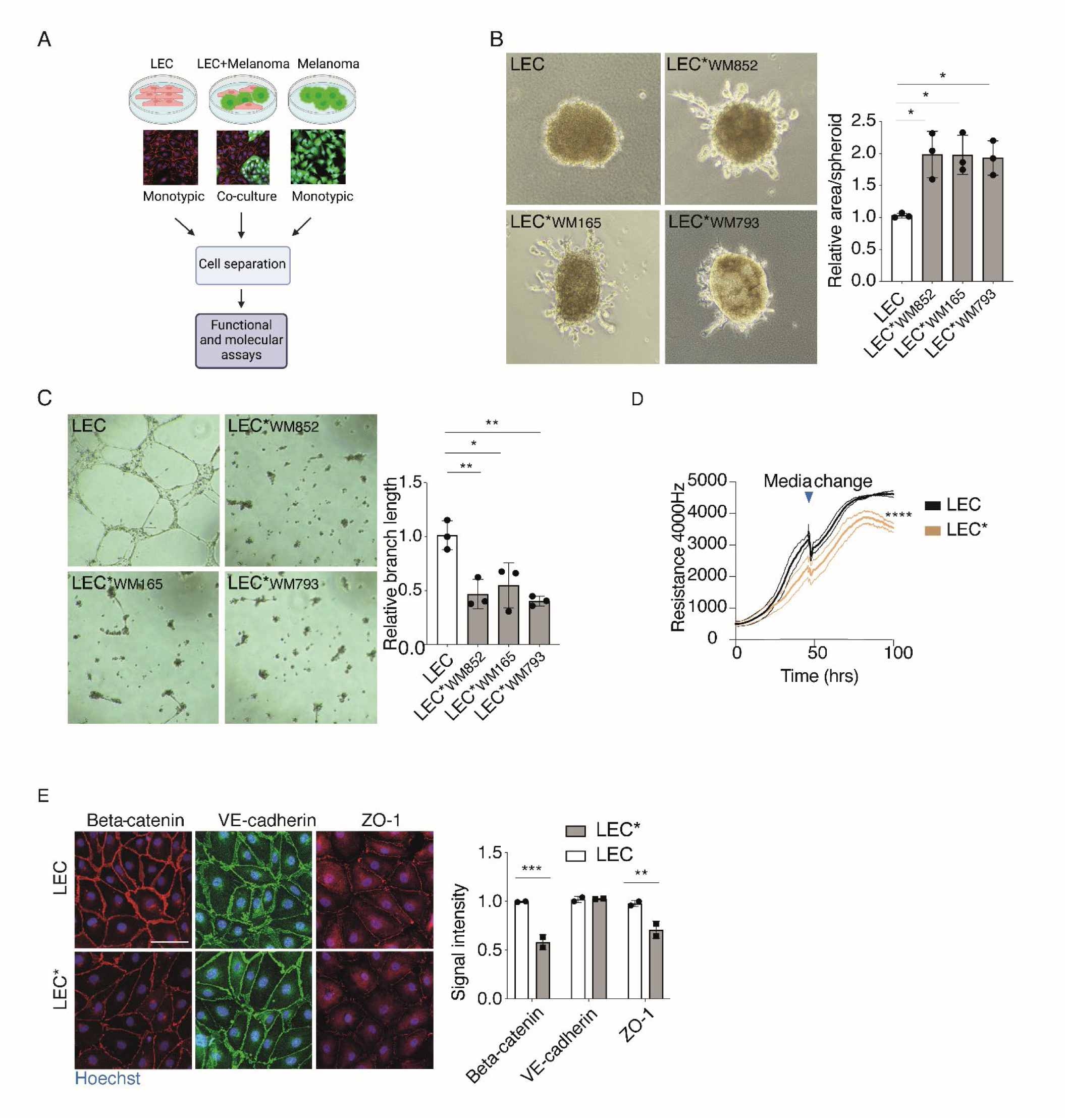
Melanoma cells induce functional changes in LECs. **A)** Schematic of the workflow for the monotypic and co-culture cell models. Representative immunofluorescent images are shown with green representing the GFP-expressing melanoma cells and red the LECs labeled with anti-CD31. Nuclei were counterstained with Hoechst 333424 (blue). Figure generated by BioRender.com. **B)** Spheroid sprouting assay of LECs cultured as a monotypic control culture (LEC) or as a melanoma cell co-culture (LEC*; name of the used melanoma cell line is indicated as a subscript) for two days before cell separation by FACs. Representative images of 30 spheroids after four days in fibrin are shown. Quantification of sprouting area from three independent experiments of at least three spheroids per condition is shown on the right panel. Bars, mean +/- SD. **C)** Tube formation assay LECs and LECs* cultured and sorted as in A and seeded onto Cultrex for 16h. Representative images from three independent experiments are shown on the left panel and quantification of branch length on the right panel. Bars, mean +/- SD. **D)** Monotypic control LECs and LECs* originating from a co-culture with WM852 melanoma cell line were subjected to an electrical cell impedance assay after two days of culture. A representative assay of two independent experiments is shown. Thicker lines indicate mean values and thinner lines +/- SD. **E)** IF images of the indicated proteins of monotypic LECs or LECs* originating from a co-culture with WM852 melanoma cell line. Nuclei were counterstained with Hoechst 33342. Representative images from two independent experiments are shown. Scale bar = 50 µm. Bars, mean +/-SD. *, P<0.05, **, P<0.01, ****, P<0.0001.

First, we utilized a spheroid sprouting assay where preformed LEC spheroids were embedded into a cross-linked 3D fibrin matrix. Fibrin was chosen since it is frequently deposited within the melanoma tumor microenvironment and perivascular niche *in vivo* (12). During the four-day incubation in 3D matrix, the monotypic control LEC spheroids remained as round spheres, while an outgrowth of sprouts was observed in the LEC* spheroids derived from melanoma cell co-cultures with all the cell lines tested (Figure 1B), indicating a clear phenotypic change in the LECs after the melanoma cell co-culture.

To determine the capacity of LECs to form capillary-like structures, we exploited a classical tube formation assay in which LECs are cultured overnight on a reconstituted basement membrane extracellular matrix support (Cultrex). Monotypic LECs spontaneously formed tube-like structures suggestive of a clear vasculogenic cell behavior, whereas the LECs* originating from melanoma cell co-cultures largely failed to form continuous tubular networks (Figure 1C).

LEC monolayer permeability as an indicator of its barrier function was addressed using non-invasive electrical cell impedance monitoring. LECs* from melanoma cell co-cultures of WM852 and WM165 showed an impaired ability to form insulating barriers compared to the control LEC layers (Figure 1D and Supplementary figure 1A). This suggests that cell-cell contacts are weakened upon co-culture with melanoma cells. Therefore, we next analyzed expression of well-known proteins at the cell-cell contact sites in the monotypic control LECs and LECs*. In line with this, after melanoma cell co-culture, significantly less beta-catenin and ZO-1 signal was detected in LECs* when compared to the parental, monotypic control LECs (Figure 1E). Although we did not see significant reduction in VE-cadherin intensity at the cell junctions, its staining pattern in LECs* was less serrated and reticular, possibly indicating weaker cell-cell junctions (13, 14). These results suggest that melanoma cell interaction with LECs may induce changes in the LEC junction maturation processes.

Lastly, we addressed the effect of co-culture on LEC proliferation (Supplementary figure 1B). After cell sorting, monotypic control LEC and LECs* co-cultured with WM852 melanoma cell line were subjected to EdU based click-it chemistry to identify proliferating cells. No significant changes in the ratio of proliferating cells were seen between the monotypic control and LECs*.

Taken together, clear phenotypic and functional changes were observed in LECs* after the co-culture with melanoma cells. The changes might at least partly be due to changes in the cell-cell contacts, as suggested by the weaker ability of the LECs* to form insulating barriers and the altered levels and distribution of the analyzed cell junctional proteins.

To uncover molecular changes in LECs induced by the melanoma cell co-culture, we performed single-cell RNA sequencing (scRNAseq) to compare the gene expression profiles of monotypic LEC (control LEC) to LECs* co-cultured with the metastatic melanoma cell line WM852. To this end, two different samples were prepared and subjected for scRNAseq analysis. First, LECs from a monotypic culture were mixed with melanoma WM852 cells (1:10 ratio of melanoma:LEC) from a monotypic culture (Sample 1, Figure 2A) in order to identify the possible residual melanoma cells originating from the co-culture samples after separation of the two cell types. For the LEC* sample, LECs were co-cultured for two days with WM852 melanoma cells after which the two cell types were separated and the LECs* subjected to further analysis (Sample 2). Upon sample analysis and clustering in UMAP plots nine different cell clusters could be identified (Figure 2A). Feature heatmaps of melanoma (SOX10) and LEC (PROX1) markers showed that melanoma cells localized to clusters 5 and 8 and the remaining clusters consisted of LECs and LECs* (Figure 2A and Supplementary figure 2A). When the distribution of LECs within the clusters was analyzed, we found that in some clusters either the LECs* (Sample 2) or control LECs (Sample 1) were heavily enriched (Figure 2B). For example, cluster #1 mainly consisted of the LECs* (Sample 2) whereas cluster #2 predominantly represented the control LECs (Sample 1). In cluster #1, genes regulating angiogenesis and cell migratory processes were upregulated (Figure 2C). When LECs were compared to LECs* within cluster #1, we found that for instance, inflammatory response pathways were also upregulated in LECs* when compared to LECs (Supplementary excel file, Supplementary figure 2B). By RT-qPCR we further validated selected genes that were found highly upregulated in the LEC* sample and involved in the vascular development (EDN1, ENG) or inflammatory response (CCL2, IL32) pathways (Supplementary figure 2C).

**Figure 2.**
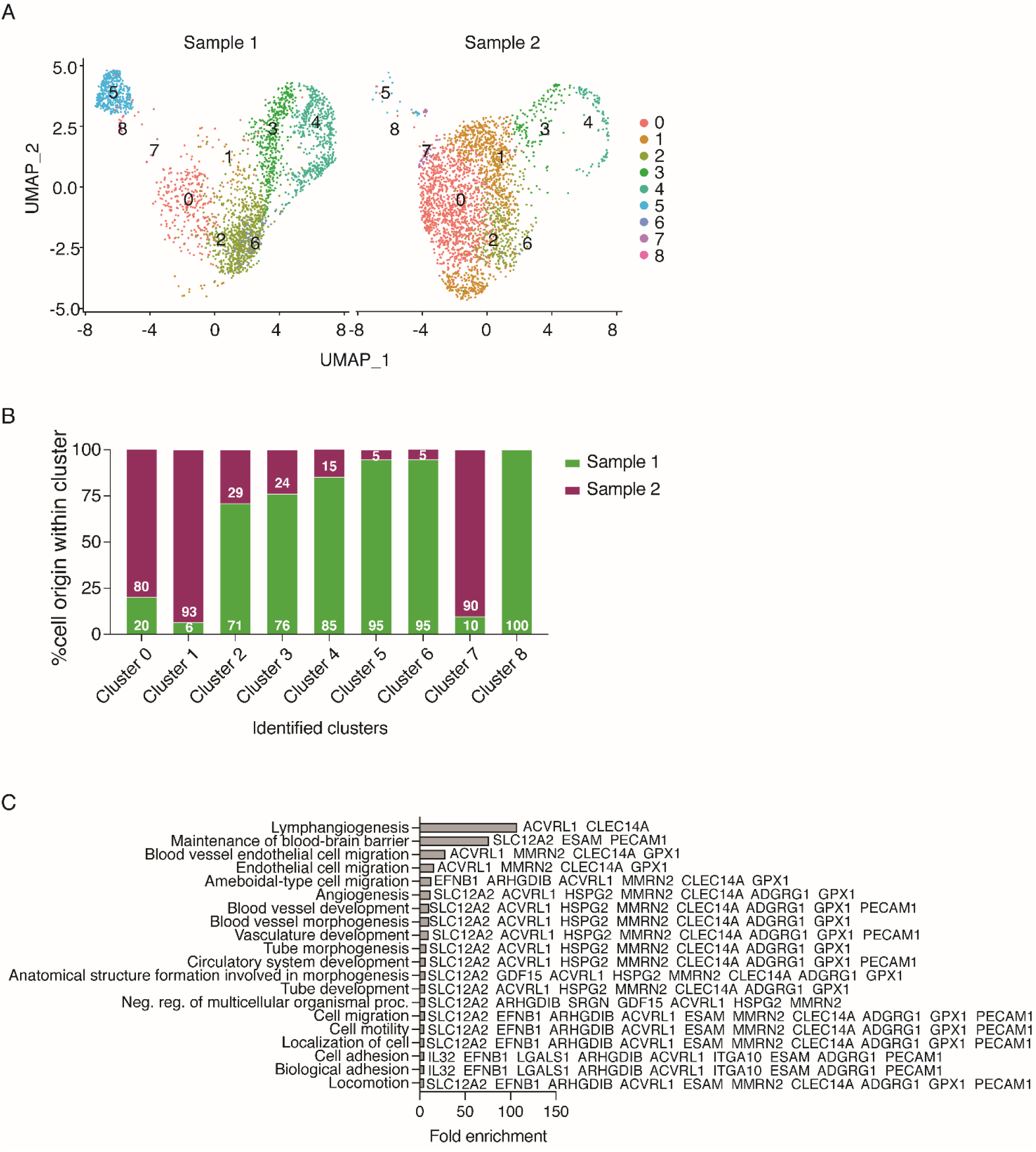
Melanoma cells induce gene expression changes in LECs. **A)** UMAP clustering plots of the cells in Sample 1, consisting of a mixture of monotypic control LECs and monotypic WM852 melanoma cells, and in Sample 2 consisting of LECs* co-cultured for two days with WM852 melanoma cells and sorted for analysis. **B)** Distribution of the cells in Samples 1 and 2 within the clusters. **C)** Pathway analysis by ShinyGO 0.77 of genes upregulated in cluster #1 (consisting mainly of co-cultured LECs) cell population. Upregulated genes involved in the pathways are shown.

These results indicate that melanoma co-culture induces phenotypic alterations in LECs including alteration of cell sprouting in 3D, angiogenic potential and barrier functions and significant changes in gene expression levels of the pathways likely regulating these processes.

### Melanoma cell derived WNT5B contributes to the functional changes in LECs

To elucidate how the melanoma cells induce the observed changes in LECs* we first addressed if they require direct cell-cell contact between the two cell types or are mediated by a paracrine factor(s). To this end, we carried out tube formation assays in which the control LECs were cultured in conditioned media (CM) derived from either the monotypic cultures of LECs or WM852 or from the WM852-LEC co-culture. The ability of LECs to form tubular structures was already disrupted when they were cultured in media originating from the monotypic melanoma cell culture. The tube formation capacity was disturbed to a greater extent with CM from the WM852-LEC co-culture (Figure 3A). This suggested that a secreted factor(s) from the melanoma cells could be responsible for causing this functional change in LECs.

**Figure 3.**
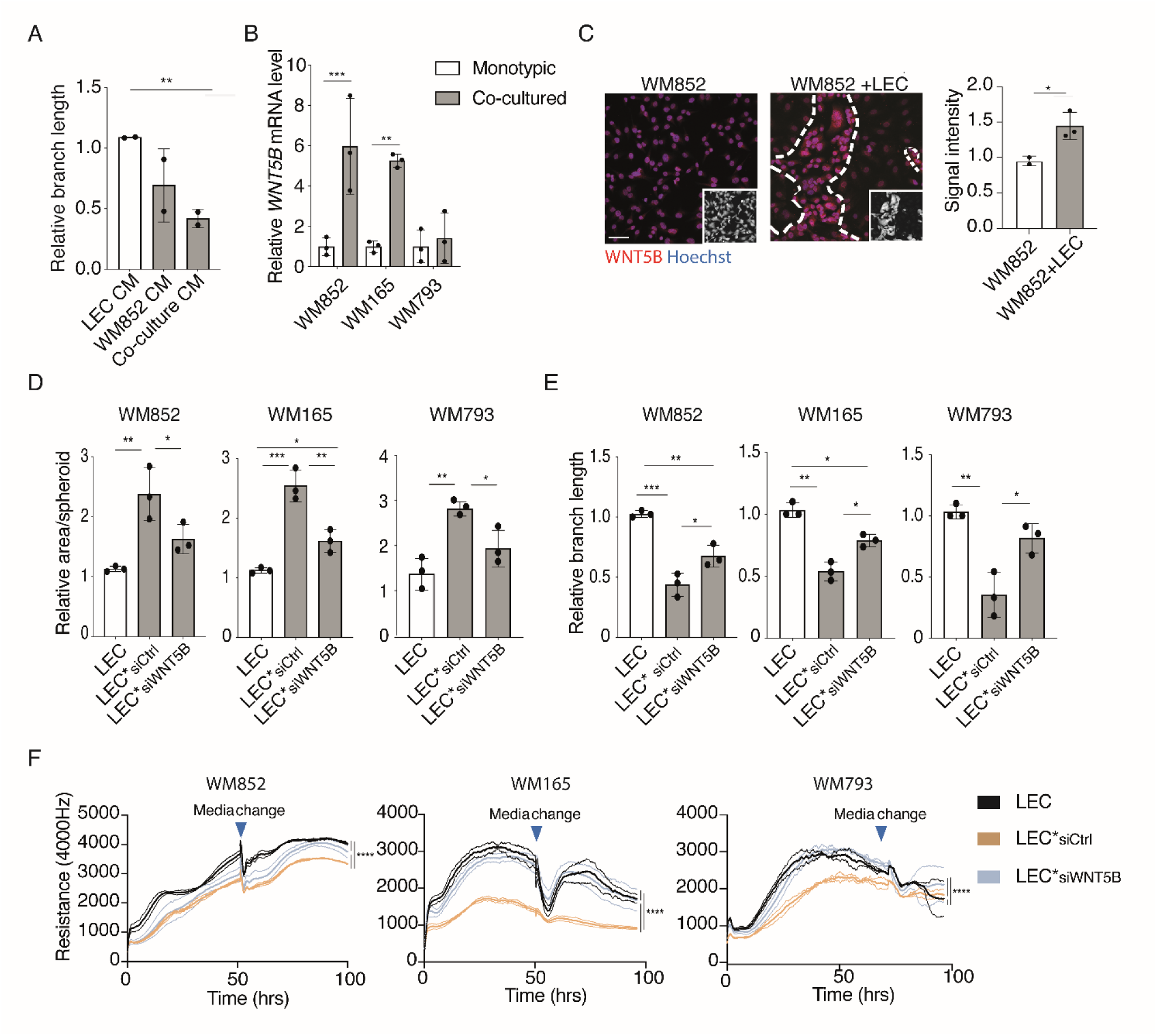
Melanoma cell derived WNT5B contributes to the functional changes in LECs. **A)** Quantification of the relative branch length of a tube formation assay with LECs cultured in conditioned medias (CM) from monotypic LEC, WM852 or LEC+WM852 co-culture for 24 h and subjected to a 16 h tube formation assay. Experiment was performed two independent times. Bars, mean +/- SD. **B)** RT-qPCR of *WNT5B* mRNA levels in the indicated monotypic or LEC co-cultured melanoma cell lines from two independent experiments. Bars, mean +/- SD. **C)** IF images of monotypic WM852 and WM852+LEC cultures labeled with an antibody against WNT5B. Small inserts identify the GFP-expressing melanoma cells. Nuclei were counterstained with Hoechst 33342. Representative images from three independent experiments are shown on the left panel, and quantification of WNT5B relative signal intensity from at least 100 cells/experiment/condition is shown on the right panel. Scale bar = 50 µm. Bars, mean +/- SD. **D)** Quantification of the spheroid sprouting assay of monotypic LEC and LECs co-cultured with the indicated melanoma cells (LEC*). Prior to the co-culture melanoma cells were pretreated with control (siCtrl) or *WNT5B* targeting siRNAs (siWNT5B) for 24h. Graph shows a mean of three independent experiments each with at least four spheroids/condition quantified. Bars, mean +/- SD. **E)** Quantification of the tube formation assay with LECs cultured and treated as in D. n=3. Bars, mean + SD. **F)** The electric cell impedance assay with LECs cultured and treated as in D. Graph indicates mean +/- SD for each sample. Representative experiment is shown. *, P<0.05, **, P<0.01, ***, P<0.001, ****, P<0.0001.

To identify the factor(s) we sought for possible candidates from our previously published RNAseq data, where gene expression profiles of melanoma cells before and after the LEC co-culture had been determined (11). In this dataset, WNT5B, a gene encoding a secreted signaling molecule, was found to be highly upregulated in the LEC co-cultured WM852 melanoma cells. The role of WNT5B in cancers has not been extensively studied, but there are indications of WNT5B having tumor promoting roles. For instance, WNT5B promotes proliferation and invasion of certain breast cancer and oral squamous cell carcinoma cell lines (15–17), in genomic analyses it associates with the most aggressive pancreatic cancer subtype (18), and pancreatic cancer cells that have undergone mesenchymal transition have been shown to promote the metastatic potential of the neighboring epithelial cells (19).

We next confirmed that *WNT5B* mRNA was upregulated upon LEC co-culture in the metastatic melanoma cell lines WM852 and WM165. We also analyzed *WNT5B* induction following LEC co-culture with a non-metastatic melanoma cell line WM793 (Figure 3B) but did not see any increase in *WNT5B* mRNA. Further analysis showed that this cell line expressed *WNT5B* at about 10-fold higher level when compared to WM852 and WM165 cell lines already before the LEC co-culture (Supplementary figure 3-1A) suggesting different, intrinsic regulation of WNT5B expression in this cell line. The upregulation of WNT5B protein in the co-cultured WM852 and WM165 cells was also confirmed by IF (Figure 3C and Supplementary figure 3-1B).

To investigate if the melanoma cell derived WNT5B could be responsible for the changes observed in the LEC* phenotype and function, melanoma cells were pretreated with siRNA targeting the *WNT5B* gene or control siRNA prior to the co-culture with LECs. Co-culture with the melanoma cells depleted of *WNT5B* (Supplementary figure 3-1C) significantly reduced the sprouting of LEC spheroids (Figure 3D). Depletion of *WNT5B* in the melanoma cells could partially restore the tube formation ability of LECs* (Figure 3E) and the barrier function of the LEC* layer (Figure 3F). Accordingly, treatment with the recombinant WNT5B protein was sufficient to reduce the barrier function of the control LEC layer (Supplementary figure 3D) and decrease beta-catenin and ZO-1 expression on the cell membrane (Supplementary figure 3-1E). Together, these functional assays demonstrate that WNT5B expressed and secreted by the melanoma cells contributes to the functional changes in LECs*.

WNT ligands signal through binding to the Frizzled receptors and additional co-receptors on the target cell membrane (20). To investigate which receptor on the LEC surface would be responsible in transmitting the WNT5B signal we first characterized the mRNA expression of the ten known human Frizzled receptors in parental, control LECs and LECs* (Supplementary figure 3-2 A). Out of these receptors, five were expressed at detectable levels in LECs. Frizzled 6 and 8 were the most abundantly expressed receptors in LECs, but we did not observe significant differences in any of the Frizzled receptor mRNAs between the control LECs and LECs*. As previously shown by molecular docking experiments, WNT5B has the highest binding affinity to Frizzled 8(21). We therefore chose to focus on Frizzled 6 and 8 in the further experiments and treated the LECs with either control siRNA or siRNAs targeting *FZD6* or *FZD8* genes for 24h before the start of the 48h co-culture with WM852 melanoma cells followed by sorting and functional assays. We did not observe any differences between the LECs* and control LECs in the spheroid sprouting assay or tube formation assays when either *FZD6* or *FZD8* was depleted (Supplementary figure 3-2 B-C). This suggests that there are other receptors or contribution by multiple receptors mediating the downstream effects of WNT5B in LECs.

### WNT5B facilitates melanoma cell escape from the primary injection site

To further characterize the role of WNT5B in melanoma progression we chose to use an *in vivo* tumor model where melanoma cells are injected intradermally into the mouse ear pinna, which is rich in lymphatic capillary networks and feasible to image by confocal microscopy. To this end, we first treated WM852 melanoma cells with control siRNA or siRNA targeting *WNT5B* prior to the 48h co-culture with LECs and subsequent cell separation (Figure 4A). 5 x 10^5^ cells from each condition were implanted in Matrigel and allowed to grow for one week after which the mice were sacrificed, and the ears processed for whole mount immunofluorescence and imaging. After seven days significantly fewer cells were seen in the ear samples injected with the LEC co-cultured melanoma cells (siCtrl*) compared to the injection sites of monotypic control melanoma cells (siCtrl) (Figure 4B). Moreover, the siCtrl* melanoma cells displayed a diffuse growth phenotype (see arrowheads in Figure 4B) when compared to the siCtrl melanoma cells (indicated with dashed line in Figure 4B), suggesting more invasive character of these LEC co-cultured cells (11). Since we had not seen significant differences in the proliferation rates between the siCtrl*and siCtrl cells *in vitro* (Supplementary figure 4A) this suggested that the siCtrl* melanoma cells might have escaped from the primary injection site via the lymphatic vasculature more efficiently than the siCtrl cells. When cells pretreated with siRNA targeting WNT5B (siWNT5B*) prior to the co-culture were injected, they were better retained at the initital injection site when compared to the siCtrl* cells (Figure 4B). Interestingly, the implanted siWNT5B* cells showed a similar, diffuse growth phenotype comparable to siCtrl* cells indicating that the melanoma derived WNT5B was primarily affecting the passage through lymphatic vasculature rather than the growth phenotype of melanoma cells.

**Figure 4.**
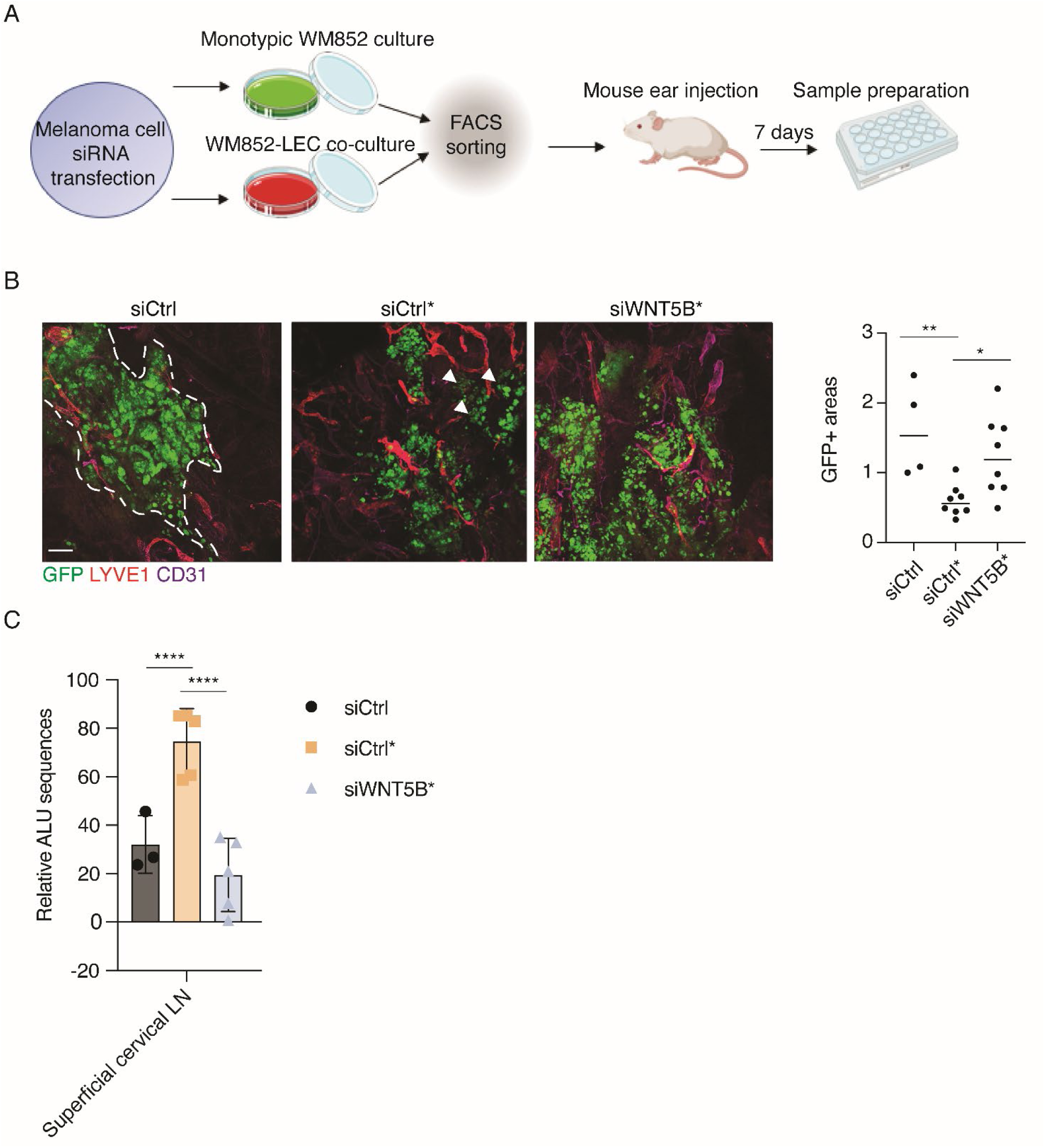
WNT5B facilitates melanoma cell escape into draining lymph nodes. **A)** Schematic of the workflow. WM852 melanoma cells were treated with siRNAs for 24 h and cultured as monotypic cultures (siCtrl) or with LEC (siCtrl*, siWNT5B*). After two days, the two cell types were separated and melanoma cells were injected intra dermally into mouse ear pinna. After one week, mice were sacrificed and ears, lungs, liver and superficial and inguinal lymph nodes were harvested and processed for analyses. Schematics generated with BioRender.com. **B)** Representative images of the GFP-expressing WM852 melanoma cells (siCtrl, siCtrl*, siWNT5B*) in mouse ear pinna epidermis. Dashed line indicates the boundaries of injected melanoma cells, arrowheads show the diffuse growth phenotype of the siCtrl* melanoma cells. The relative size of areas occupied by GFP-expressing melanoma cells was quantified from each mouse ear. Relative size for the GFP positive area of each mouse ear is shown. (siCtrl, n=4; siCtrl* n=8; siWNT5B*n=8, n refers to number of ears quantified). Scale bar = 200µm. Bars, mean +/- SD. **C)** qPCR for the relative human ALU sequences from the mouse superficial cervical lymph nodes. Mouse genomic actin was used as a control. Single values for each mouse are shown, (siCtrl, n=3; siCtrl* n=5; siWNT5B*n=5). Bars, mean +/- SD. *, P<0.05, **, P<0.01, ****, P<0.0001.

We next hypothesized that the siCtrl* cells would first locally invade into the lymphatic capillaries in the ear, and further drain into the cervical sentinel lymph nodes in the mouse neck. To assess this, we harvested superficial cervical and inguinal lymph nodes and quantified the human Alu sequences, as an indication of the presence of human melanoma cells in the lymph nodes (Figure 4C and Supplementary figure 4B). As a negative control we used mouse inguinal lymph nodes (Supplementary figure 4B), since we did not expect that the short duration of the experiment would be sufficient for melanoma cells to metastasize from the initial injection site to more distant lymph nodes. The relative Alu sequence amount was significantly higher in the superficial cervical lymph nodes collected from the mice implanted with siCtrl* melanoma cells when compared to the siCtrl ones, supporting more efficient escape of the siCtrl* cells from the primary injection site (Figure 4C). Importantly, less Alu sequence signal was found in the superficial cervical lymph nodes of the mice implanted with siWNT5B* cells when compared to mice implanted with siCtrl* cells. These data demonstrate that WNT5B plays an important role in promoting melanoma metastasis *in vivo* from the primary injection site to the sentinel lymph nodes, most probably through its effects on the LECs.

### Notch3 regulates WNT5B expression in melanoma cells

We have previously shown that Notch3 is highly upregulated in melanoma cells upon co-culture with LECs and is important for melanoma invasion and metastasis (11). Interestingly, Notch3 binding to the WNT5B promoter area has been previously reported, but not validated, in ovarian cancer cells (22). Therefore, we tested if Notch3 would function as the upstream regulator of WNT5B expression in the co-cultured melanoma cells. To this end, we set up co-cultures of LECs with WM852 and WM165 melanoma cells pretreated with control siRNAs (siCtrl*) or siRNAs targeting *NOTCH3* 24 h prior to the start of the 48 h co-culture. The siCtrl treated monotypic melanoma cells were used as a control. When compared to the siCtrl* cells the *WNT5B* mRNA was significantly reduced in the siNOTCH3* pretreated cell lines (Figure 5A and Supplementary figure 5A), suggesting that Notch3 was activating *WNT5B* transcription. *NOTCH3* depletion in the monotypic melanoma cells, however, did not significantly affect the basal level of *WNT5B* (Figure 5A and Supplementary figure 5A). To confirm the role of Notch3 as an inducer of *WNT5B* expression, the co-culture assays were also carried out in the presence of the Notch pathway inhibitor DAPT or vehicle as a control (EtOH). When compared to vehicle treated co-cultures the DAPT treatment significantly reduced the upregulation of *WNT5B* mRNA levels in WM852 and WM165 cells co-cultured with LECs (Figure 5B and Supplementary figure 5B), further supporting a key role for Notch3 in regulating WNT5B expression in the metastatic WM852 and WM165 melanoma cells. However, the non-metastatic melanoma cell line WM793, with endogenously high *WNT5B* levels, did not show any significant change in the *WNT5B* mRNA levels upon siNOTCH3 or DAPT treatments, suggesting other than Notch3 regulatory pathways for the sustained, high WNT5B expression in this cell line (Figure 5A-B and Supplementary figure 5A-B).

**Figure 5.**
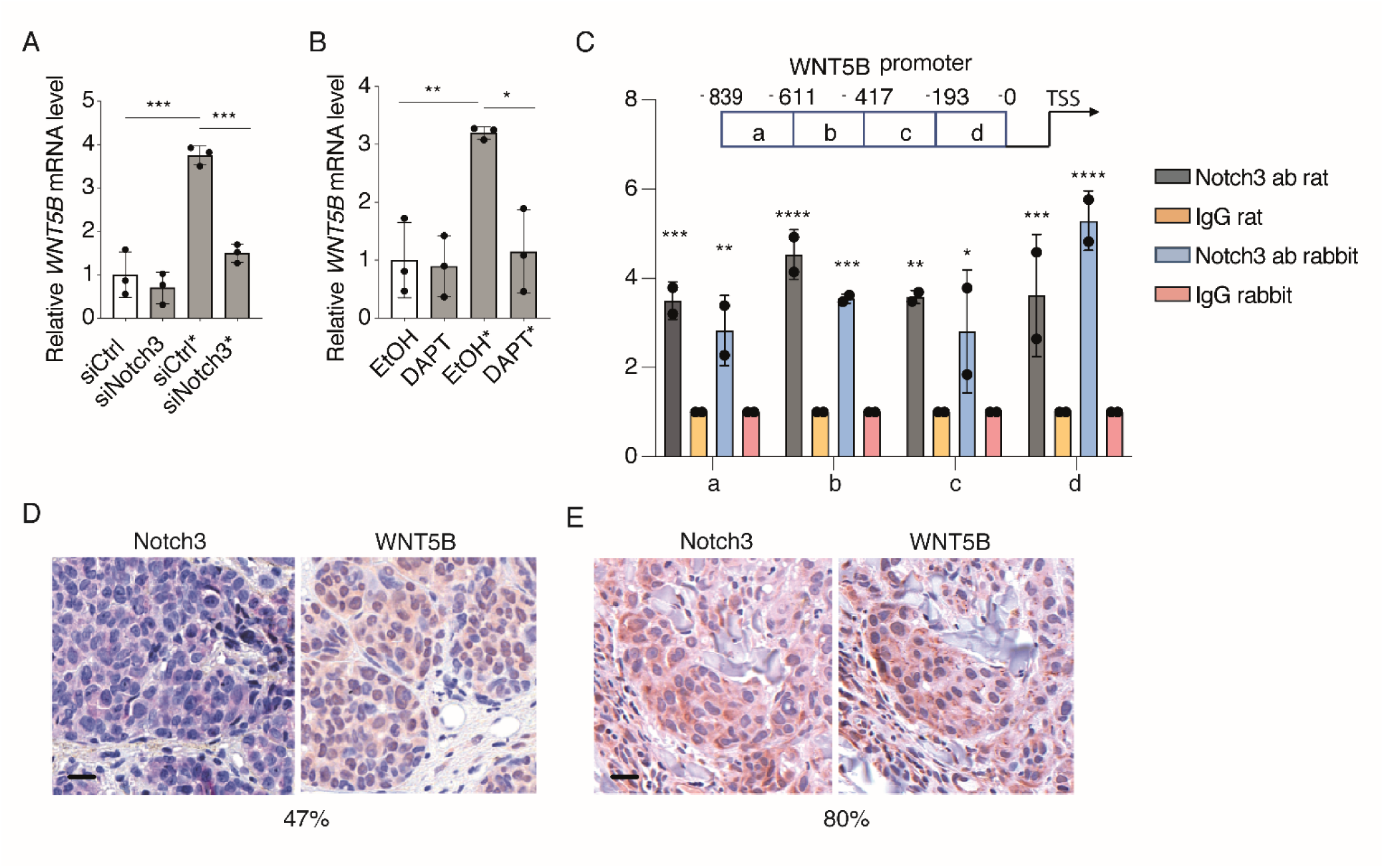
Notch3 regulates WNT5B expression in melanoma cells. **A)** WM852 cells were pretreated with the indicated siRNAs and subjected to monotypic or LEC co-cultures (*). After cell sorting, *WNT5B* mRNA levels in melanoma cells were measured by RT-qPCR. Graph shows results from three independent experiments. Bars, mean +/- SD. **B)** WM852 cells were cultured as monotypic cultures or in co-culture with LECs (*) and treated with vehicle (EtOH) or DAPT. WNT5B mRNA level was measured in the sorted melanoma cells by RT-qPCR. n=3. Bars, mean +/- SD. **C)** Top panel; schematic presentation of the *WNT5B* promoter area amplified by qPCR following ChIP. Numbers indicate the nucleotides upstream of the *WNT5B* transcription start site. Bottom panel; ChIP from WM852 cells expressing ectopic NICD3 for 24 hours. ChIP was performed with two different anti-Notch3 antibodies and respective control IgGs and DNA was amplified from the indicated promoter regions of the *WNT5B* gene. Bars, mean + /- SD. **D-E)** Representative images of human primary melanoma tumor (D) and metastasis samples (E) labeled for the indicated proteins. Percentage below the images indicates the proportion of samples with Notch3 and WNT5B signal co-localization. Scale bar = 20 µm. *, P<0.05, **, P<0.01, ***, P<0.001, ****, P<0.0001.

To confirm the role of Notch3 as a transcriptional activator of *WNT5B*, we next addressed the binding of Notch3 on the *WNT5B* promoter region by ChIP-PCR. To this end, WM852 melanoma cells were first transfected with a plasmid encoding an active Notch3 intracellular domain, NICD3, and subjected to ChIP on day two using two different anti-Notch3 antibodies. As shown in Figure 5C, Notch3 binds to the *WNT5B* gene promoter region thus confirming its role as a transcriptional activator of WNT5B in melanoma cells.

We next investigated if Notch3 and WNT5B expression would correlate in melanoma patient samples (Table 1). We chose to use a patient cohort, where, despite a confirmed negative sentinel lymph node biopsy (SNB) at the time of diagnosis, the patients later presented with metastases. At the time of the diagnosis, 36 out of 55 patients had tumors less than 4 mm deep measured by the Breslow thickness, thus representing patients to whom the sentinel lymph node involvement is routinely used as a diagnostic tool. This cohort was also atypical since patients’ age at the time of diagnosis did not correlate with survival time. Their disease progressed very aggressively, with an average survival of 3.75 years (Supplementary figure 5C), in comparison to an average 5-year survival of 99.5 % for a patient with localized melanoma (https://seer.cancer.gov/statfacts/html/melan.html). Primary tumor samples from 55 melanoma patients and 45 metastasis samples were stained with antibodies against WNT5B and Notch3. 46 and 47 samples, respectively, were successfully scored of the primary tumors and 41 from the metastases. Co-distribution of expression of the two proteins was found in 47 % of the primary tumor samples (Figure 5D) and in 80 % of the samples from the first site of metastasis (Figure 5E), providing further support for the importance of the Notch3-WNT5B axis in melanoma and its relevance in the aggressive melanoma cases.

Together these data demonstrate that Notch3 acts as an upstream positive regulator of WNT5B in the metastatic melanoma cell lines WM852 and WM165. Furthermore, we found co-distribution of Notch3 and WNT5B expression in the primary and metastatic samples from a cohort of melanoma patients with exceptionally aggressive disease.

### Notch ligand DLL4 is a potent inducer of Notch3 and WNT5B in melanoma

We next set to investigate how Notch3 activation is induced in the melanoma cells upon interaction with LECs in co-culture. Notch signaling is initiated when the Notch receptor binds to its ligand, a transmembrane protein on the cell membrane of a neighboring cell. This induces a pulling force to the Notch receptor thus exposing the two cleavage sites of the receptor for proteases. The cleavage allows the release of the intracellular domain of the receptor (NICD) and its nuclear translocation where it binds to additional cofactors to initiate target gene transcription (23). Human cells express all of the five Notch ligands, Dll1, Dll3-4 and Jag1-2 (24). We therefore first addressed which of these ligands are expressed in LECs and LECs*. RT-qPCR analysis showed that they both express all five Notch ligands at detectable levels and no significant differences were observed between the monotypic LEC and co-cultured LECs*(Supplementary figure 6A). Of all the ligands, JAG1 and JAG2 showed the highest mRNA expression levels in LECs. To test which ligand on the LEC membrane mediates activation of Notch3 on the melanoma cells, we cultured different melanoma cell lines on cell culture dishes coated with recombinant, immobilized Fc-fragments of all the Notch ligands to mimic the pulling force of the ligand on the Notch receptor. After two days, cells were harvested and their mRNA levels of *NOTCH3* as well as Notch3 downstream targets *HEY1* and *HES1* were measured (Figure 6A). DLL4-Fc induced the expression of *NOTCH3* mRNA in the metastatic melanoma cell lines WM852 and WM165, while DLL1 induced *NOTCH3* only in WM165. Of the known Notch downstream targets *HEY1* was upregulated in WM852 and WM165 by DLL4-Fc and to some extent also by DLL1-Fc albeit not significantly. We did not observe any clear changes in the expression level of *HES1*. The other tested Notch ligands did not induce any significant changes in *NOTCH3* or the downstream targets.

**Figure 6.**
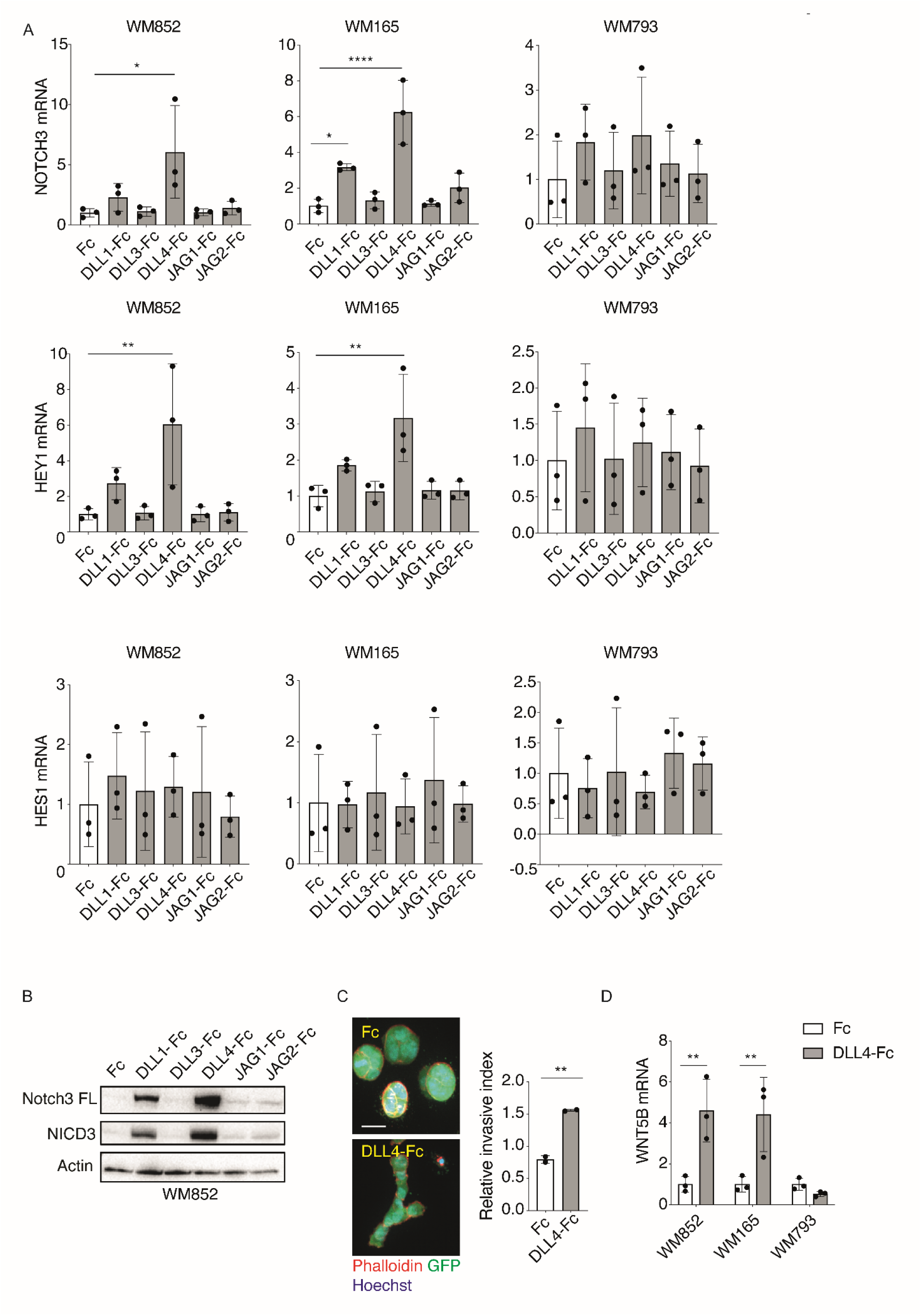
Notch ligand DLL4 is a potent inducer of Notch3 and WNT5B in melanoma. **A)** Indicated melanoma cell lines cultured on dishes precoated with the indicated Fc fusion fragments of Notch ligands or Fc as a control for two days, and subjected to RT-qPCR of *NOTCH3*, *HEY1* and *HES1*. n=3. Bars, mean +/- SD. **B)** Immunoblotting of WM852 cells cultured as in A for the indicated targets (FL= full length). A representative blot of three independent experiments is shown. **C)** A 3D fibrin droplet invasion assay of WM852 cells cultured on DLL4-Fc and Fc as in A. GFP expressing melanoma cells were stained with Phalloidin 594 and nuclei were counterstained with Hoechst 33342. Maximum intensity of Z-projections of the confocal stacks are shown. Graph shows quantification of the relative invasive index from two independent experiments with at least 50 cell clusters quantified/condition. Scale bar = 20µm. Bars, mean +/- SD. **D)** RT-qPCR of *WNT5B* levels in melanoma cell lines cultured as in C. Experiment was performed three independent times. Bars, mean +/- SD. *, P<0.05, **, P<0.01, ***, P<0.001.

We also assessed the activation of Notch3 by immunoblotting. Culturing of the melanoma cell lines on the DLL4-Fc coated dishes induced an increase of the cleaved, active NICD3, especially in the WM852 and WM165 and to a lesser extent in WM793 (Figure 6B and Supplementary figure 6B-C). NICD3 induction by DLL1-Fc coating was also seen in WM852 and WM165 but it was less pronounced compared to the DLL4-Fc mediated induction. Interestingly, the full length Notch3 receptor level was also increased by DLL1-Fc and DLL4-Fc treatments, which together with the observed increase in the mRNA levels (Figure 6A) suggest that Notch3 is induced through an autoregulatory loop in the melanoma cells.

Our previous report demonstrated that the LEC-induced Notch3 is crucial for the increased melanoma invasion (11). We next addressed the ability of DLL4-Fc to increase melanoma invasion by embedding the ligand activated melanoma cells into a cross-linked 3D fibrin matrix for four days. According to our earlier results, DLL4-Fc could significantly increase the invasive growth of WM852 and WM165 melanoma cells into the surrounding fibrin (shown as invasive index in Figure 6C and Supplementary figure 6D).

Lastly, we measured the *WNT5B* expression levels in the cells cultured on the Fc-fragments of Notch ligands by RT-qPCR. DLL4-Fc coating induced about a four-fold increase in *WNT5B* expression in WM852 and WM165 (Figure 6D). In WM165 also DLL1 and JAG2 induced about a two-fold increase in the *WNT5B* level, although it did not reach significance (Supplementary figure 6E). In line with the earlier results, no significant changes in *WNT5B* expression were observed upon culturing the WM793 cells on top of the five different Notch ligand Fcs (Figure 6D and Supplementary figure 6E).

These results show that DLL4 is a potent inducer on the LEC surface activating the Notch3 receptor on the melanoma cells upon co-culture, leading to increased invasion and upregulation of *WNT5B* in the metastatic melanoma cell lines.

## DISCUSSION

Lymphatic endothelium has recently gained attention as an active, cancer promoting element with functions beyond simply acting as a passive route for cancer cell dissemination. Here, we report a novel DLL4-Notch3-WNT5B signaling axis induced upon melanoma cell and LEC interaction and promoting melanoma metastasis via its effects on LEC phenotype and function. Specifically, we show that DLL4, expressed on the LEC surface, activates Notch3 in melanoma cells, which in turn triggers *WNT5B* transcription and protein expression in melanoma cells. The melanoma derived WNT5B then contributes to the functional changes in LECs in a paracrine manner (Figure 7).

**Figure 7.**
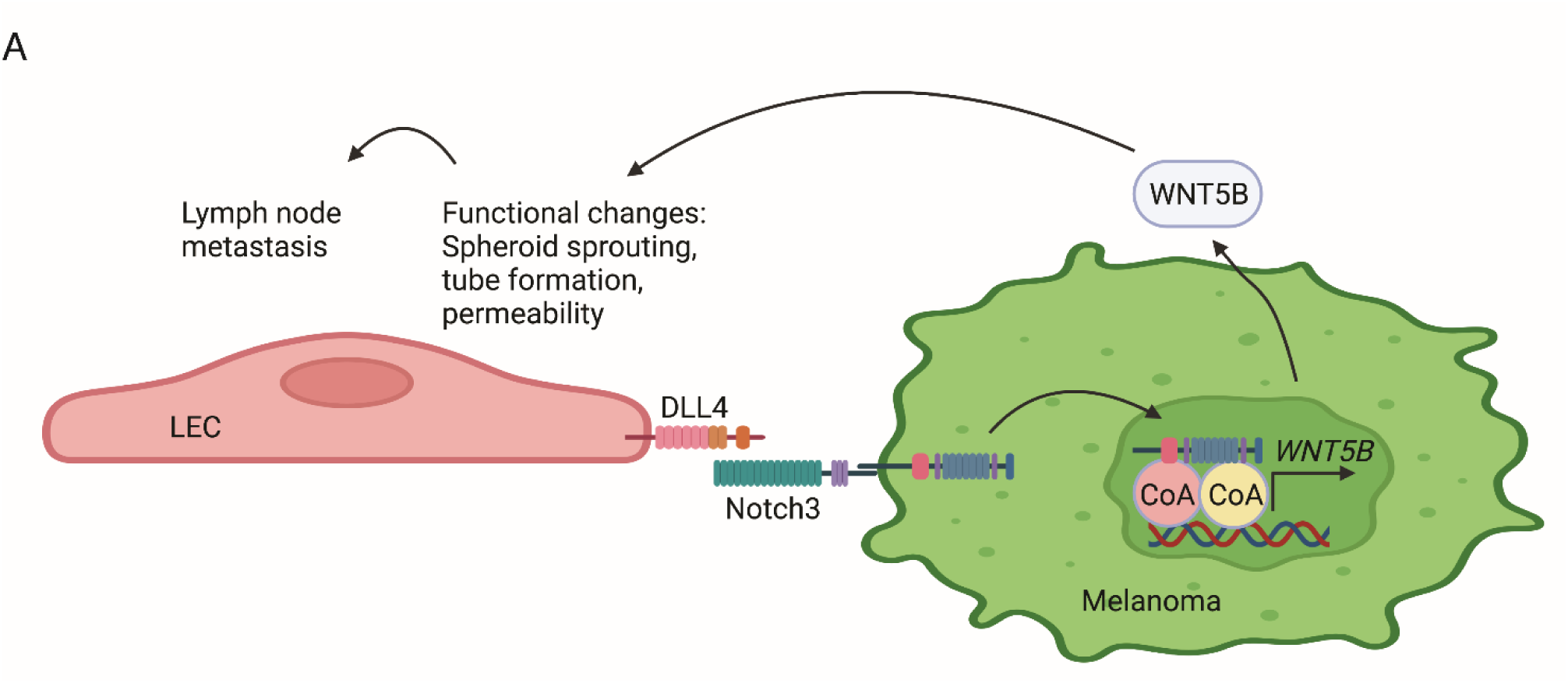
Model of the bi-directional melanoma-LEC crosstalk. **A)** Schematic model of the bi-directional melanoma cell crosstalk with LECs and the role of Notch3 in the LEC functional changes through induction of WNT5B.

Following the melanoma co-culture, we found upregulation of genes associated with inflammatory responses in the LECs*. These transcriptional changes are likely to contribute to the disruption of lymphatic junctions thus leading to increased permeability of LEC monolayers and facilitating cancer dissemination. LECs can act both as the target of inflammatory signals and represent a remarkable source of tumor promoting cytokines. As an example, it has been shown that tumor exposed LECs significantly increase their IL6 production, and thereby promote tumor cell invasion and proliferation (25). Our scRNAseq data is in line with a previous study in mice where they found that pathways regulating immunomodulatory responses were upregulated in tumor draining lymph node LECs compared to control naïve lymph node LECs (26). Our scRNAseq data revealed upregulation of genes involved in vascular developmental processes such as Endothelin-1 and Endoglin (27). Endothelin-1 produced by ECs has been shown to promote melanoma cell migration and vessel-like channel formation, and, interestingly, LECs cultured in the media containing Endothelin-1 also show enhanced cell migration (28). Endoglin expression increases in ECs during vascular damage (29) and inflammation and it facilitates the infiltration of inflammatory cells into the endothelium (30, 31). All these molecular changes along with others found in our scRNAseq data likely contribute to the tumor promoting, phenotypic changes in LECs.

We identified WNT5B as a novel key component of the bi-directional, reciprocal melanoma-LEC crosstalk. WNT5B upregulation was mediated by Notch3 activation induced in melanoma cells by direct interaction with LECs upon co-culture. Moreover, our results demonstrate that the melanoma derived WNT5B participates in inducing the functional changes in LECs in a paracrine manner. The mouse ear xenograft assay indicated that WNT5B facilitates the LEC primed melanoma cell escape from the primary injection site and translocation to the proximate draining lymph nodes. Although WNT5B clearly contributes to melanoma cell escape from the initial injection site, other factors are also likely to contribute to this process; as an example, we found that the LEC primed melanoma cells appear to grow as a more dispersed cell colony when compared to the monotypic cells, possibly generating more opportunities for melanoma cells to communicate with the mouse lymphatic vasculature and promote lymphatic dissemination.

WNT5B, a member of the WNT family proteins, has been described as an inducer of the non-canonical, beta-catenin independent WNT signaling pathways and is often considered as an antagonist of the canonical WNT signaling. It has been reported to regulate cell migration, proliferation, and differentiation during development (32). WNT5B is upregulated in triple negative breast cancer patient samples and correlates with a worse prognosis (15). It is also associated with lung adenocarcinoma where its expression correlates with lymph node metastasis and poor survival (33). To our knowledge the role of WNT5B in melanoma metastasis has not been characterized before. Interestingly, in oral squamous cell carcinoma, WNT5B has been shown to be upregulated in tumor cells when compared to normal tissue (17). In addition, WNT5B was shown to induce partial endothelial-to-mesenchymal transition (End-MT) in LECs by increasing alpha-SMA expression and reducing VE-cadherin expression. Moreover, WNT5B depletion was shown to reduce lymph node metastases of oral squamous cell carcinoma cells; suggesting that WNT5B could be more broadly utilized by cancers to facilitate lymph node metastases (17). In melanoma mouse models it has been shown that metastasized melanoma cells in lymph nodes can invade into the lymph node associated blood vessels and disseminate directly to distant organs (34, 35). It is therefore possible that WNT5B is one of the key players contributing to the first metastatic steps in melanoma progression, and our results imply that melanoma cells require a direct interaction to LECs in the tumors for the initiation of the metastatic cascade.

We further found that Notch ligand DLL4, expressed in LECs, is a potent inducer amongst the tested ligands of Notch3 in melanoma cells. Notch3 is a central protein in altering the melanoma cell behavior; its effects include induction of increased cell migration (36), stem-like cell characteristics (37) and invasion and metastasis (11). We and others have previously demonstrated that endothelial cells induce Notch3 expression in melanoma cells upon interaction (11, 36, 37). Here we show that Notch3 is not only contributing to the more aggressive melanoma cell characteristics but can also elicit functions that modify the melanoma tumor microenvironment, and thereby promote cancer progression. Accumulating evidence supports the contribution of Notch3 to the aggressiveness of melanoma. Therefore, it represents an appealing target for aggressive melanoma therapies. However, until now the clinical trials using Notch pathway inhibitors have not been successful, mainly due to the unspecific nature of currently available Notch inhibitors, which often lead to severe gastrointestinal side effects. Therefore, a deeper knowledge of the upstream and downstream effects of Notch3 signaling in melanoma may provide better cues for targetable molecular candidates. For instance, several phase I trials have been launched where Porcupine inhibitors were used to abolish WNT secretion from cancer cells, aiming at targeting cancer cell proliferation and cancer stem cells (38). These inhibitors might represent viable therapeutic modalities in targeting the tumor microenvironment as well.

Our study and increasing number of other evidence demonstrate that instead of simply acting as a route for metastatic and inflammatory cell transport, the lymphatic system is an active player shaping the cancer cell behavior, which in turn the cancer cells can actively manipulate to further promote tumorigenesis. Therefore, the bidirectional crosstalk and interaction between the tumor and lymphatic system can potentially provide novel molecular targets and opportunities to target cancer cell invasion and metastasis through combinatorial therapies.

## METHODS

### Cell lines

Primary human dermal lymphatic endothelial cells (LECs) were obtained from Lonza and cultured in endothelial growth medium (EBM-2, Lonza) supplemented with growth factors included in the supplement bullet kit (except for the VEGF supplement) and 5% fetal calf serum (FBS; full media referred as EGM-2). EGM-2 media was also used to grow LEC-melanoma co-cultures. WM852, WM165 and WM793 (Wistar Institute, Philadelphia, PA) were cultured in Dulbecco’s Modified Eagle Medium (DMEM, Lonza) supplemented with 10% FBS and 1% penicillin/ streptomycin. The cells were transduced with dual eGFP-luc (pMX-Rg) or GFP reporters (pLENTI6-eGFP, Invitrogen) as described in (39). All cultures were grown in standard conditions (37°C, 5% CO2) and regularly tested negative for mycoplasma using Eurofins Mycoplasmacheck service (https://eurofinsgenomics.eu/en/genotyping-gene-expression/applied-genomics-services/mycoplasmacheck/). The identity of the melanoma cell lines was authenticated by the FIMM (university of Helsinki) cell line authentication service (https://www.helsinki.fi/en/infrastructures/genome-analysis/infrastructures/fimm-genomics/cell-line-authentication).

### RNA interference and cell culture treatments

Cells were cultured to approximately 60% confluency on 24-or 6-well plate and transfected with siRNAs at final concentrations of 10-25nM using lipofectamine RNAiMAX (Invitrogen) according to manufacturer’s instructions. Cells were cultured in the presence of siRNA for 24h before using in subsequent assays. The following siRNAs were used: non-targeting control (Ambion, MA4390843; Dharmacon, D-001810-10-05) Notch3 (Ambion, 4392420; Dharmacon, L-011093-00-0005) WNT5B (Dharmacon, L-009761-00-0005), FZD6 (L-005505-00-0005, Dharmacon) and FZD8 (L-003962-00-0005, Dharmacon).

Where indicated, gamma-secretase inhibitor DAPT (Sigma) at 10µM concentration and recombinant WNT5B protein (7347, R&D Biosystems) at 1000ng/ml were applied.

### Transient transfection

WM852 cells were seeded on 10 cm cell culture dishes one day prior transfection to reach 80-90% confluency next day. Cells were then transfected with 8.8 μg of NICD3-pCLE (Addgene) using Fugene HD (Promega) according to manufacturer’s instruction. Next day, the cells were used for subsequent analysis.

### Notch ligand coating

24 or 12 well plates were coated with the Fc domain, as a control, or Notch ligand fused with the Fc domain at a concentration of 10µg/ml for 6 h at room temperature (RT) (DLL1 Fc, 10184-DL-050 Biotechne; DLL3 Fc, DL3-H5255 ACRO Biosystems; DLL4 Fc, 158-10171-H02H-100 Sino Biological; JAG1 Fc, 158-11648-H02H Sino Biological, and JAG2 Fc, 1726-JG-050 R&D Systems). 0.4 - 0.75*10^5 cells were cultured on coated plates for 48h, then harvested and used for subsequent assays.

### Melanoma cell-LEC co-cultures and cell sorting

For the co-culture of LECs with melanoma cells, cells were seeded on fibronectin (Sigma) coated cell culture dishes in 1:2-3:4 melanoma-LEC ratio in EGM-2 media. For the first experiments, the cell separations were performed using magnetic separation of LECs and melanoma. For this separation process, melanoma cells were first loaded with dextran coated nanoparticles at 1mg/ml (fluid-MAG-DX, Chemicell) for 24 h. Dextran-loaded melanoma cells were cultured with LECs for 48 h after which the cells were sorted using MidiMacs separator and LS columns (Miltenyi Biotec). Due to the cessation of the production of these magnetic nanoparticles, we had to change the separation to be carried out by FACS (Sony SH800Z). The sorted cells were used for the subsequent functional or molecular assays.

### 3D fibrin assays

The fibrin assays were adapted from a previously described angiogenesis assay (40), described in detail in (41). To study the melanoma cell invasion in 3D fibrin, 5000 melanoma cells were embedded in crosslinked fibrin (Calbiochem) and cultured for 72h. In the LEC spheroid assay, 5,000 LECs were first seeded on a U-bottom 96-well plate pre-coated with 1% agarose (Invitrogen) o/n to let the LECs form spheroids under non-adherent conditions. Next day the spheroids were harvested and embedded in a fibrin matrix. Four days later, the fibrin droplets containing either the LEC spheroid or melanoma cells were fixed with 4% PFA and stained as indicated for subsequent analysis. The droplets were imaged using either the Zeiss LSM780 confocal microscope or Nikon Eclipse TS2 phase contrast microscope and the images were analyzed with ImageJ software.

### Tube formation assay

10^4 LECs were seeded on 96-well plates pre-coated with 50 μl of Cultrex (3433, R&D Systems) and incubated 24 h in the cell incubator. Phase contrast images of the cells, four fields per well, were obtained using Nikon Eclipse TS2 phase contrast microscope and the tubular structures were quantified with ImageJ software.

### Electric cell-substrate impedance assay

2*10^4 LECs were seeded onto 10 mM L-cysteine and 5μg/ml fibronectin-coated 96W10idf 96-well plate (Ibidi) and the resistance of the cell monolayers was recorded at 4000 Hz by an ECIS Z Theta instrument connected to a computer running an ECIS software version 1.4.8 (Ibidi) over four days. The media was changed once during that time. The readout indicates the changes in resistance over time.

### RNA isolation and Real-time quantitative PCR (qRT-PCR)

NucleoSpin II kit (Macherey Nagel) was used to isolate RNA according to manufacturer’s instructions. 200-1000ng of RNA was reverse transcribed with the Taqman reverse transcription kit (N8080234, Applied Biosystems). The transcript levels of *NOTCH3* were measured using QuantiTect primer assay (QT00003374, Qiagen), the sequences of the other targets are provided in Table 2. The reactions were performed using SYBRGreen PCR mix (Applied Biosystems).

### Western blot

Western blot analysis was performed as previously described (11). Cells were lysed in RIPA buffer supplemented with protease and phosphatase inhibitors (Thermo Scientific). The protein amount of the lysates was measured with Bio-Rad protein assay dye reagent concentrate (Bio-Rad). Equal amounts of protein were loaded on 4-15% SDS-PAGE gel (Bio-Rad) and the gel was run at 55 mA for 50 min. Proteins were transferred onto nitrocellulose membrane (Bio-Rad) using Bio-Rad Trans-Blot Turbo and the membrane was blocked using 5% non-fat dry milk in TBST (Tris-buffered saline with 0.1% Tween 20). The probing was done either for 1h at RT or o/n at +4°C using following antibodies: rabbit anti-Notch3 (M-143, Santa Cruz) or mouse anti-β-actin (A1978, Sigma). The primary antibody incubation was followed by incubation in HRP-conjugated secondary antibody for 1h RT at room temperature (goat anti mouse IgG and goat anti-rabbit IgG, Cell Signaling Technology). Bands were detected by chemiluminescence using ECL solution (WesternBright Sirius, Advansta) and visualized by Chemi-Doc (Bio-Rad).

### Immunofluorescence staining

2D cell cultures were labelled as previously described (11) with the antibodies against WNT5B (ab94914, Abcam), beta-catenin (610153, BD Biosciences), ZO-1 (40-2200, Invitrogen) and VE-Cadherin (555661, BD Biosciences). Secondary antibodies conjugated with Alexa594 or Alexa 647 (Invitrogen) were used to visualize the labelled cells and the nuclei were counterstained with Hoechst 33342 (Fluka). The fluorescence images were acquired using confocal LSM780 Zeiss microscope and sum of pixel intensity values was quantified using CellProfiler (Broad Institute) and normalized by number of the cells.

### Chromatin-immunoprecipitation and PCR

WM852 cells were transfected with the expression plasmid NICD3-pCLE (11) and 48h later, the chromatin was processed by the simpleChIP kit according to the manufacturer’s instructions (Cell Signaling Technology). The chromatin was precipitated using antibodies against Notch3 (M-134 Santa Cruz Biotechnology or 8G5 Cell Signaling) or normal goat or rat IgG antibody (Santa Cruz Biotechnology). DNA was purified using the PCR Purification kit (Macherey-Nagel) according to the manufacturer’s instructions.

After DNA purification, the WNT5B promoter regions were amplified and analyzed by qPCR (Table 2).

### Mouse ear xenograft assay

The NOD.CgPrkdcscid Il2rgtm1Wjl/SzJ strain female mice (age 20 to 24 weeks old) were either purchased from Scanbur or obtained as a generous gift from Timo Otonkoski (University of Helsinki, Finland). Mice were housed in a temperature controlled and pathogen-free facility with *ad libitum* access to food and water. All the animal experiments were approved by the Committee for Animal Experiments of the District of Southern Finland (license: ESAVI/10548/2019).

5*10^5 cells melanoma cells were mixed in 1:1 PBS-Matrigel and injected intradermally into the mouse ear pinna of the mice. Tumors were allowed to grow for seven days after which the mice were sacrificed. The collected ears were split into ventral and dorsal halves and fixed in 4% PFA for 25 min RT. Tumors were imaged with Zeiss LSM 780 confocal microscopy. The total size of areas covered by GFP expressing melanoma cells were quantified by ImageJ.

Superficial cervical and inguinal lymph nodes were harvested, and tissue DNA was isolated using the NucleoSpin Tissue kit (Macherey-Nagel). Presence of human melanoma cells in the collected mouse tissues was detected by quantitative PCR for the human Alu sequences while the sequences of mouse genomic actin was used as control (Table 2).

### Patient samples and Immunohistochemistry

Paraffin-embedded melanoma sections were provided by the Helsinki Biobank. All patients have given their written informed consent to the Helsinki Biobank. The study protocol and the use of the material was approved by The Ethics Committee of the Hospital District of Southwest Finland (HUS/206/2022).

Tissue sections were deparaffinized and rehydrated. The EnVision FLEX+ kit (Dako, Agilent) was used to stain the tissues. Epitope retrieval was performed using the EnVision FLEX Target Retrieval solution, high pH for 15 min at 95°C. Endogenous peroxidase blocking was performed according to the kit manufacturer. Tissue blocking was performed using the normal horse serum blocking solution (S2000, Vector Laboratories) diluted 1:20 in a Dako antibody diluent (S2022, Dako).

Sections were stained with either anti-Notch3 (1:800, HPA044392, Sigma) or anti-WNT5B antibodies (1:600, ab94914, Abcam) diluted in the Dako antibody diluent for 16h at 4°C. Secondary antibody incubation was performed according to the instructions of the kit. Chromogen reaction was performed using the Romulin AEC Chromogen kit (RAEC810, Biocare Medical) and hematoxylin staining using Mayer’s Hematoxylin (S3309, Dako). Tissues were dehydrated and mounted on coverslips using Eukitt Quick-hardening mounting medium (03989, Sigma-Aldrich). The slides were imaged with Pannoramic 250 Flash II (3DHISTEC Ltd.).

### Single-cell RNA sequencing

LECs* sorted from WM852 melanoma co-cultures (Sample 2) were mixed with control cells consisting of LECs and WM852 melanoma cells, both from monotypic cultures, in a ratio of 10:1 (Sample 1), from three replicates. The monotypic LEC and WM852 cells were mixed to identify the potential, residual melanoma cells remaining in Sample 2 after separation using magnetic beads of the co-cultured LECs and WM852. The cells were processed according to the 10X Genomics Guidelines (https://www.10xgenomics.com/support/single-cell-gene-expression/documentation/steps/sample-prep/single-cell-protocols-cell-preparation-guide). The single cell 3’ Reagent kit v2 (10X Genomics) was used to label each cell and transcript with unique molecular identifier (UMI) and to generate the cDNA library. Sequencing was performed using Illumina NovaSeq6000 sequencer. Data postprocessing and quality control were performed with the 10X Genomics Cell Ranger (version 2.1.1.) software.

The filtered count matrices were pre-processed and clustered using the Seurat V4.3.0 tool in an RStudio V2023.03.0 environment (R V4.2.3). Briefly, the filtered count matrices were imported using the *Read10X*-function and transformed into Seurat objects, filtering out cells with less than 200 features and features The filtered count matrices were pre-processed and clustered using the Seurat V4.3.0 tool in an RStudio V2023.03.0 environment (R V4.2.3). Briefly, the filtered count matrices were imported using the *Read10X*-function and transformed into Seurat objects, filtering out cells with less than 200 features and features encountered in less than three cells. Percentages of mitochondrial (pt.mito) and ribosomal RNAs (pt.rRNA) were calculated and cells were further filtered accordingly: *1)* control data (CTRL): 2000 < nFeature_RNA < 6000, pt.mito < 10, 3 < pt.rRNA < 40, and *2)* co-cultured sample data (CC): 1000 < nFeature_RNA < 4500, pt.mito < 10, 3 < pt.rRNA < 45. The final Seurat objects comprised 53.9% (2792 cells, CTRL) and 30.5% (4970, CC) of the raw cell counts. Next, the data were normalized on a logarithmic scale of 10,000, and the 2000 most variable features were retrieved using the *FindVariableFeatures*-function applying the method of variance stabilizing transformation. Data anchors were calculated between the CTRL and CC samples using *FindIntegrationAnchors* and 50 dimensions, whereafter they were integrated using the *IntegrateData*-function. The data were scaled regressing for RNA counts (nCount_RNA), pt.mito, and pt.rRNA, and principal component analysis was performed using the integrated assay. Clusters were calculated using 30 dimensions for the *FindNeighbors*-function and a resolution of 0.6 for *FindClusters*, which resulted in 9 final clusters denoted from cluster 0 to 8. Using the RNA assay, cluster markers were calculated based on the CTRL dataset using the *FindMarkers*-function with a minimum detection rate of 0.25.

The integrated Seurat object was subsetted into CTRL and CC samples, and the individual objects were further subsampled to have equal cell numbers according to the smallest number of cells encountered in the CTRL sample (2700 cells). Then, the subsampled objects were merged back together using the *merge*-function. Differentially expressed genes (DEGs) were calculated by comparing the subsampled CC and CTRL data using the *FindMarkers*-function, with a log fold change threshold set to 0.25. Upregulated genes in the CC versus CTRL samples with a *p*-adjusted value below 0.05 were used for gene ontology (GO) analysis using the ShinyGO V0.77 web platform (http://bioinformatics.sdstate.edu/go/). Here, default parameters were used to calculate the top 20 GO Biological process related to the gene lists.

### Statistical analysis

Data were analyzed with Prism 9 (GraphPad) The graphs show the mean across the biological replicates and the error bars indicate SD. The number of experimental groups is indicated in figure legends. One-way ANOVA or two-tailed t test were performed to assess the statistical significance of the differences between samples. P values lower than 0.05 were considered significant.

The ECIS data was analyzed by calculating the Area under the curve (AUC) using Prism 9. One-way ANOVA or two-tailed t test were performed to assess the significance of the differences between AUCs.

The expression levels of the proteins of interest in IHC samples were categorized into low (or negative) and high intensity groups for statistical analysis. Survival was estimated using the Kaplan-Meier method with Log-rank testing. Correlation coefficient between the markers was calculated using the Chi-square test with Phi and Cramer’s V significance testing assess the significance of the correlation using the IBM SPSS v27 software.

### Study approval

Paraffin-embedded melanoma sections were provided by the Helsinki Biobank. All patients have given their written informed consent to the Helsinki Biobank. The study protocol and the use of the material was approved by The Ethics Committee of the Hospital District of Southwest Finland (HUS/206/2022).

All the animal experiments were approved by the Committee for Animal Experiments of the District of Southern Finland (license: ESAVI/10548/2019).

## AUTHOR CONTRIBUTIONS

SA, SG and PMO conceptualized the study. SA and EM performed experiments. SJ and AM designed and collected the patient cohort and curated the patient samples and data as dermatopathologists. SA, JJ and EM performed formal analysis. SA, MHL and SK performed analysis of the scRNAseq data. OC, PS and PMO provided resources. AP, SK, KV provided guidance for experiments. SG and PMO supervised the study. PMO provided funding. SA and PMO wrote the original draft of the manuscript.

## ACKNOWLEDGEMENTS

We thank Nadezhda Zinovkina, Julia Härme, Satu Hänninen and Noora Andersson for excellent technical assistance. FIMM Institute for Molecular Medicine Finland is acknowledged for performing single cell RNA sequencing. Biomedicum Imaging Unit and Genome Biology Unit, University of Helsinki provided expert imaging services. We are grateful to the HiLife Flow Cytometry Unit, University of Helsinki, for the flow cytometry services. We thank HiLIFE Laboratory Animal Centre Core Facility, University of Helsinki, Finland for support in mouse work. We are grateful for Timo Otonkoski for providing mice for our studies.

This study was supported by the Finnish Cancer Foundation (P.M.O.) and Sigrid Juselius Foundation (P.M.O.). S.A. was supported by Doctoral Programme in Biomedicine, University of Helsinki and fellowships from Instrumentarium Science Foundation, The Finnish Cultural Foundation, Magnus Ehrnrooth Foundation, K Albin Johansson Foundation and Cancer Foundation Finland. J.J. was supported by fellowships from The Finnish Cultural foundation, Emil Aaltonen Foundation and Magnus Ehrnrooth Foundation. P.S. was supported by European Research Council (ERC) under the EU Horizon 2020 research and innovation programme (grant agreement 773076, PS), Sigrid Jusélius Foundation, Cancer Foundation Finland and Academy of Finland. M.H.L was supported by Nylands Nation. S.K. was supported by the Academy of Finland (grants 330053 and 336126).

These additional funders had no role in the study design, data collection, interpretation, or decision to submit the work for publication.

