## Supplementary Figures for "DLL4-Notch3-WNT5B axis is a novel mediator of bi-directional pro-metastatic crosstalk between melanoma and lymphatic endothelial cells"

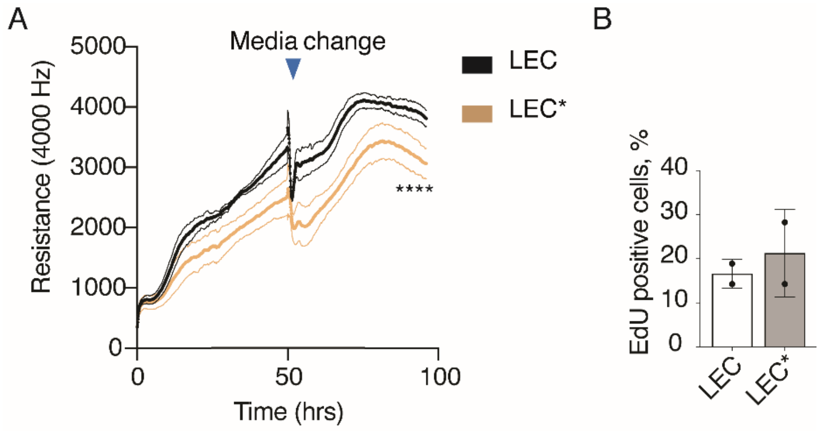


**Supplementary figure 1. Melanoma cells induce functional changes in LEC.**

**A)** Monotypic control LECs and LECs* originating from a co-culture with WM165 melanoma cell line were subjected to an electrical cell impedance assay after two days of co-culture. A representative assay is shown. Graph indicates mean +/- SD. **B)** Monotypic control LECs and LECs* originating from a co-culture with WM852 melanoma cell line were upon cell sorting treated with EdU to identify the nuclei of proliferating cells. Nuclei were counterstained with Hoechst 333424. Graph shows results from two independent experiments. Bars, mean +/- SD. ****, P<0.0001.

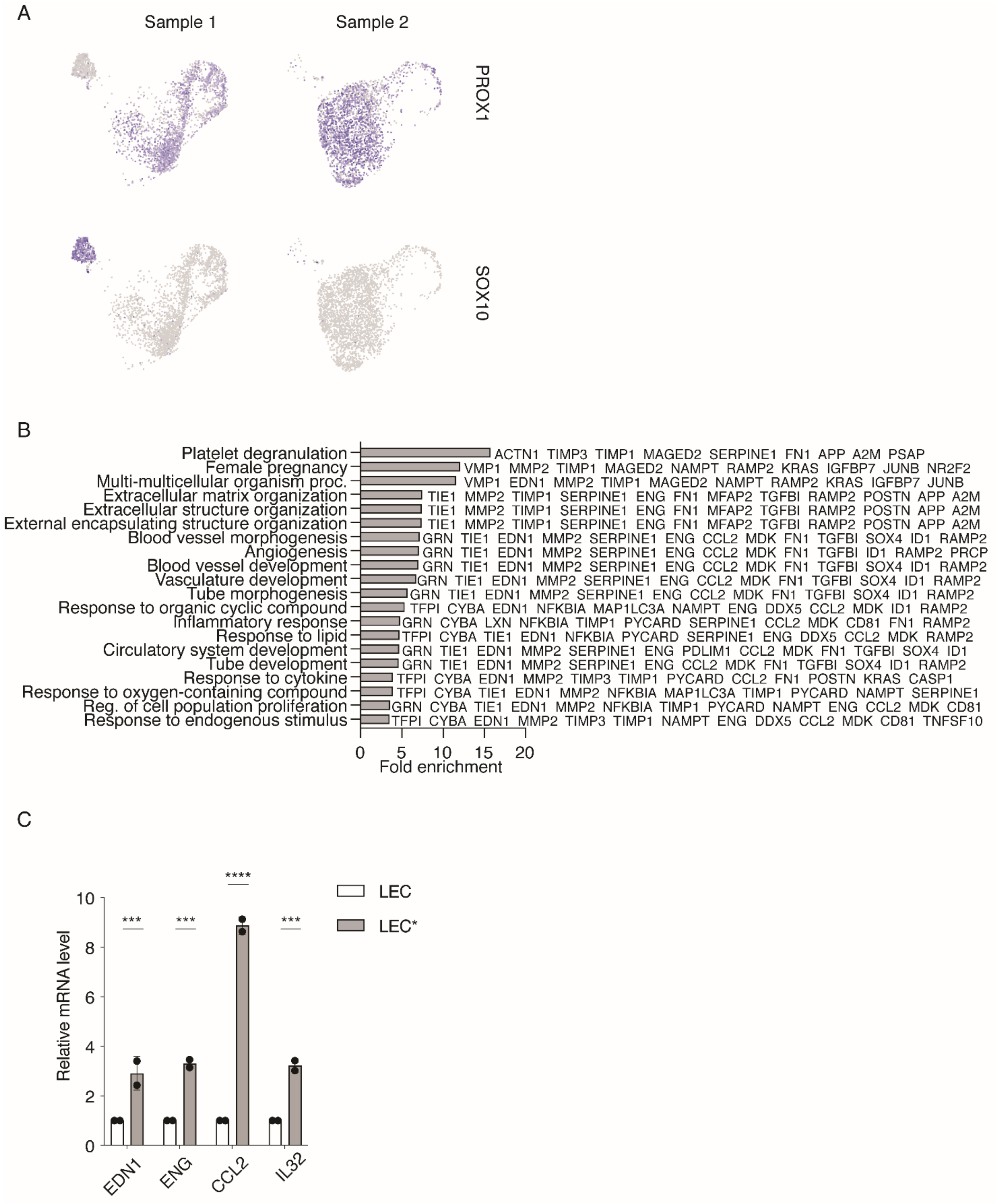


**Supplementary figure 2. Gene expression changes in LEC*.**

**A)** Feature scRNAseq heatmap visualizing the expression of SOX10 (melanoma marker) and PROX1 (LEC marker) in the cells of Sample 1, consisting of a mixture of monotypic control LECs and monotypic WM852 melanoma cells, and of Sample 2 consisting of LECs* co-cultured for two days with WM852 melanoma cells and sorted for analysis.. **B)** Pathway analysis by ShinyGo of genes upregulated in LECs* compared to control LEC within cluster #1. Selection of the genes upregulated and involved in the indicated pathways are shown. C) qPCR analysis of indicated targets in LEC* and LEC. Bars, mean +/-SD. ***, P<0.001.

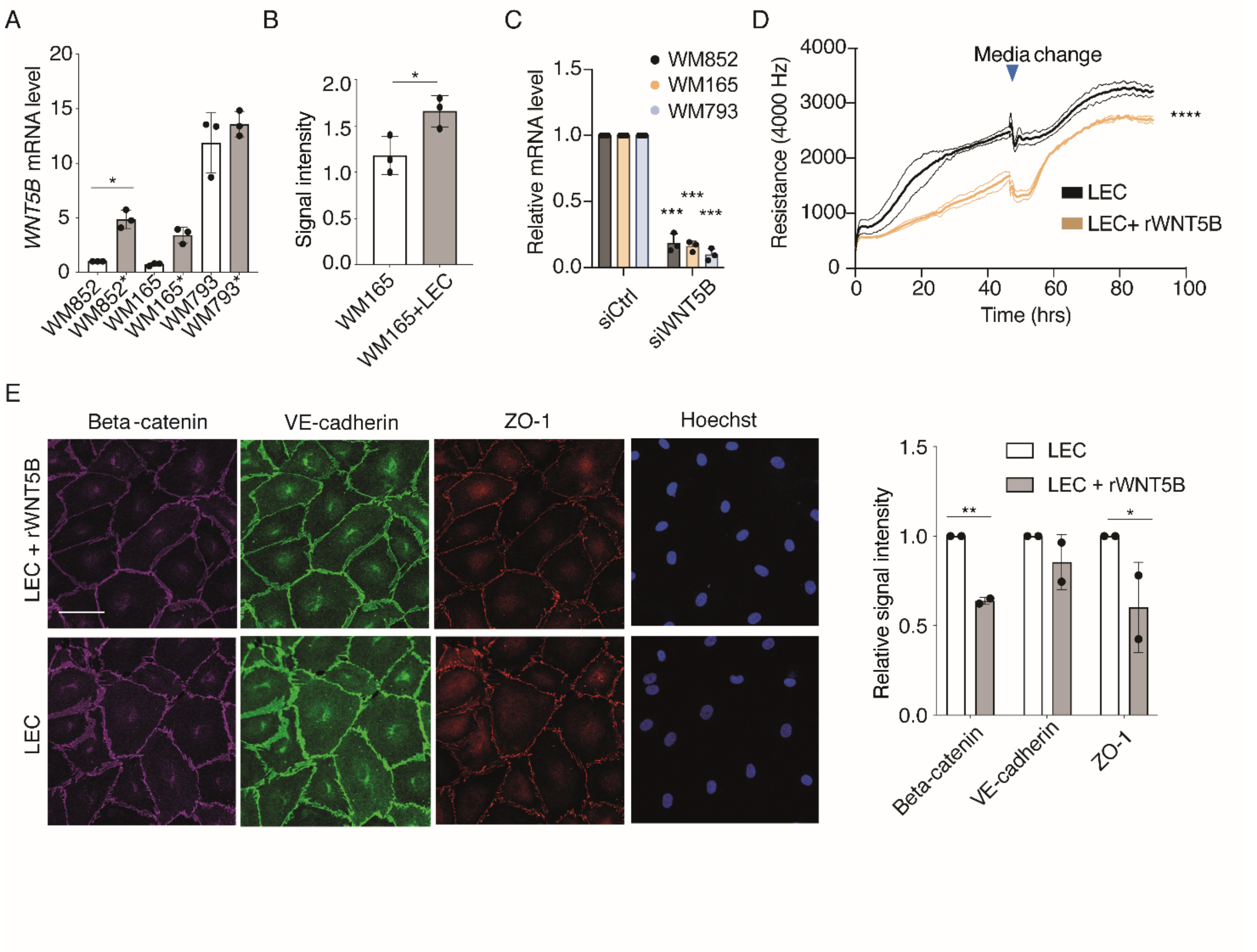


**Supplementary figure 3-1. WNT5B contributes to functional changes in LECs.**

**A)** RT-qPCR of *WNT5B* mRNA levels in monotypic and LEC co-cultured melanoma cells (*). The experiment was done three independent times. Bars, mean +/- SD. **B)** Quantification of the relative WNT5B intensity in monotypic WM165 and WM165+LEC cultures. n=3, at least 100 cells were quantified/experiment/condition. Bars, mean +/- SD. **C)** Relative *WNT5B* mRNA expression in the indicated melanoma cells treated with either control or *WNT5B* targeting siRNA for 24h before using the cells for subsequent assays. n=3. Bars, +/- SD, **D)** LECs treated with or without recombinant WNT5B at concentration of 1000 ng/ml were analyzed in an electric cell impedance assay. A representative experiment of two independent replicates is shown. Graphs indicate mean +/- SD. **E)** Control LECs or LECs supplemented with 1000ng/ml recombinant WNT5B were cultured for 16 h before fixed and labeled with antibodies for the indicated proteins. Representative images of two independent experiment is shown, bars, mean +/- SD. Scale bar = 50µm. *, P<0.05, ***, P<0.001, ****, P<0.0001.

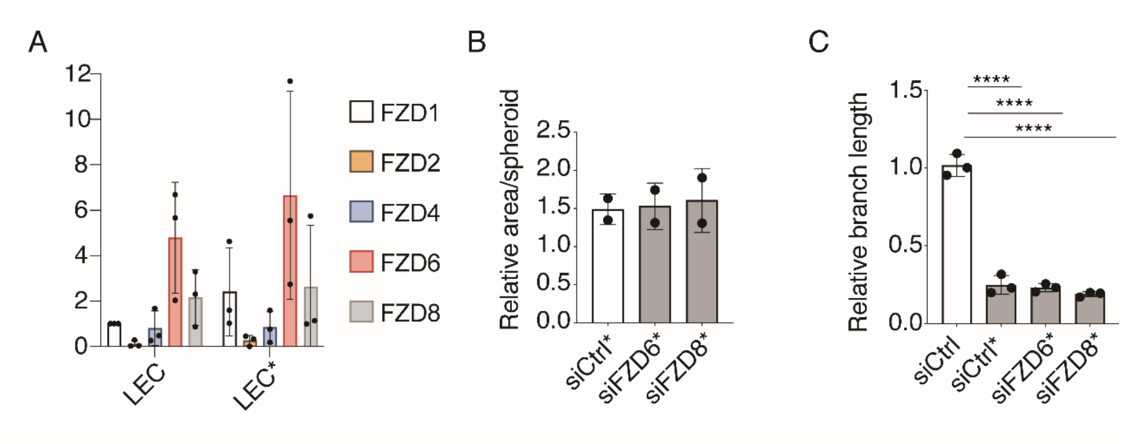


**Supplementary figure 3-2. WNT receptors FZD6 or FZD8 are not contributing to the functional changes in LEC*.**

**A)** RT-qPCR analysis of the indicated targets in monotypic LECs and LECs co-cultured with WM852 cell line (LEC*). n=3. Bars, mean +/-SD. **B)** Spheroid sprouting assay with monotypic and WM852 co-cultured LECs*. LECs were pretreated with the indicated siRNAs for one day before the start of the co-culture. Graph indicates results from two independent experiments, each with at least six spheroids quantified/condition. **C)** Tube formation assays with LECs cultured as in B. n=3. Bars, mean +/- SD. Bars, mean +/- SD. ****, P<0.0001.

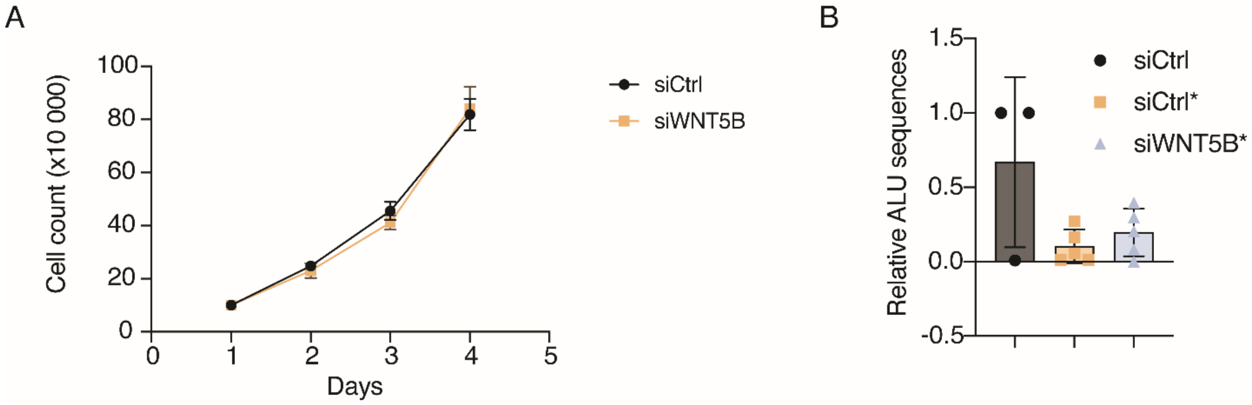


**Supplementary Figure 4. WNT5B depletion does not affect proliferation of melanoma cells.**

**A)** WM852 melanoma cell count at different time points following treatment with the indicated siRNAs. Graph shows an average of two independent experiments, error bars indicate SD. **B)** qPCR for the relative human ALU sequences from the mouse inguinal lymph nodes. Mouse genomic actin was used as a control. Single values for each mouse are shown, (siCtrl, n=3; siCtrl* n=5; siWNT5B*n=5). Bars, mean +/- SD.

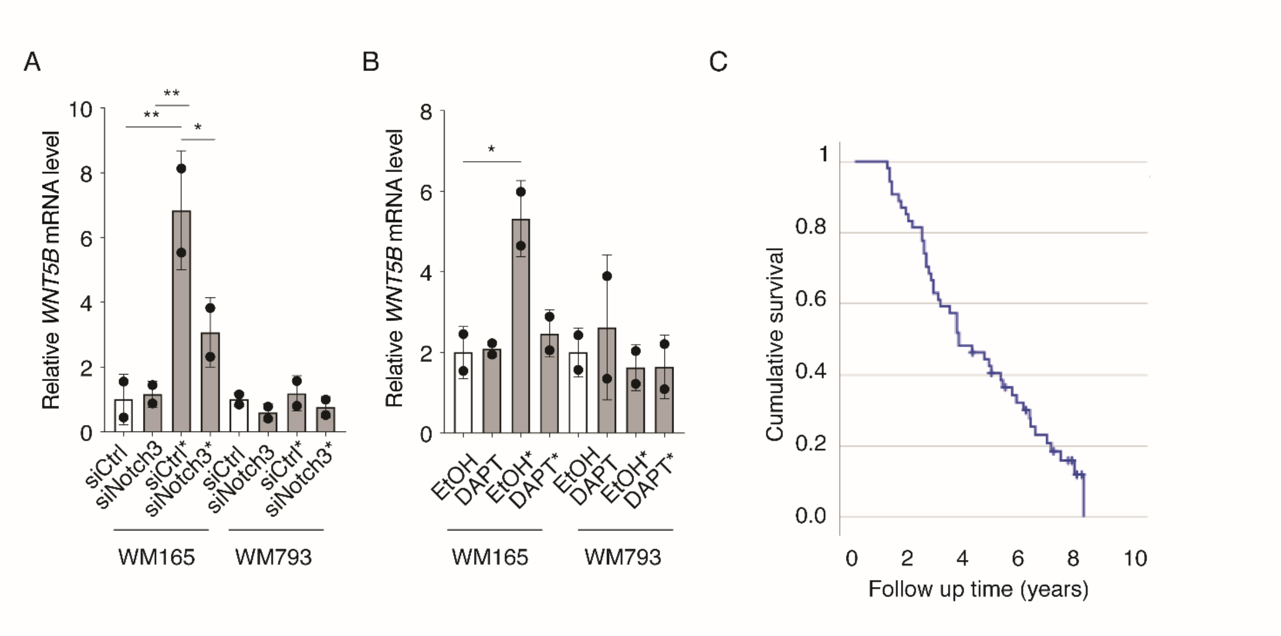


**Supplementary figure 5. Notch3 regulates WNT5B expression in melanoma.**

**A)** WM165 and WM793 cells were pretreated with the indicated siRNAs and subjected for monotypic or LEC co-culture (*). After two days, cells were sorted by FACS and *WNT5B* mRNA was measured by RT-qPCR in melanoma cells. Graph shows results from two biological replicates. Bars, mean +/- SD. **B)** WM165 and WM793 cells were cultured in monotypic cultures or in co-culture with LECs (*). The cultures were treated with either vehicle (EtOH) or with DAPT and the sorted melanoma cells were analyzed by RT-qPCR for *WNT5B* mRNA. Experiment was done two independent times. Bars, mean +/- SD. **C)** Kaplan-Meier curve of the melanoma patient overall survival. *, P<0.05, **, P<0.01.

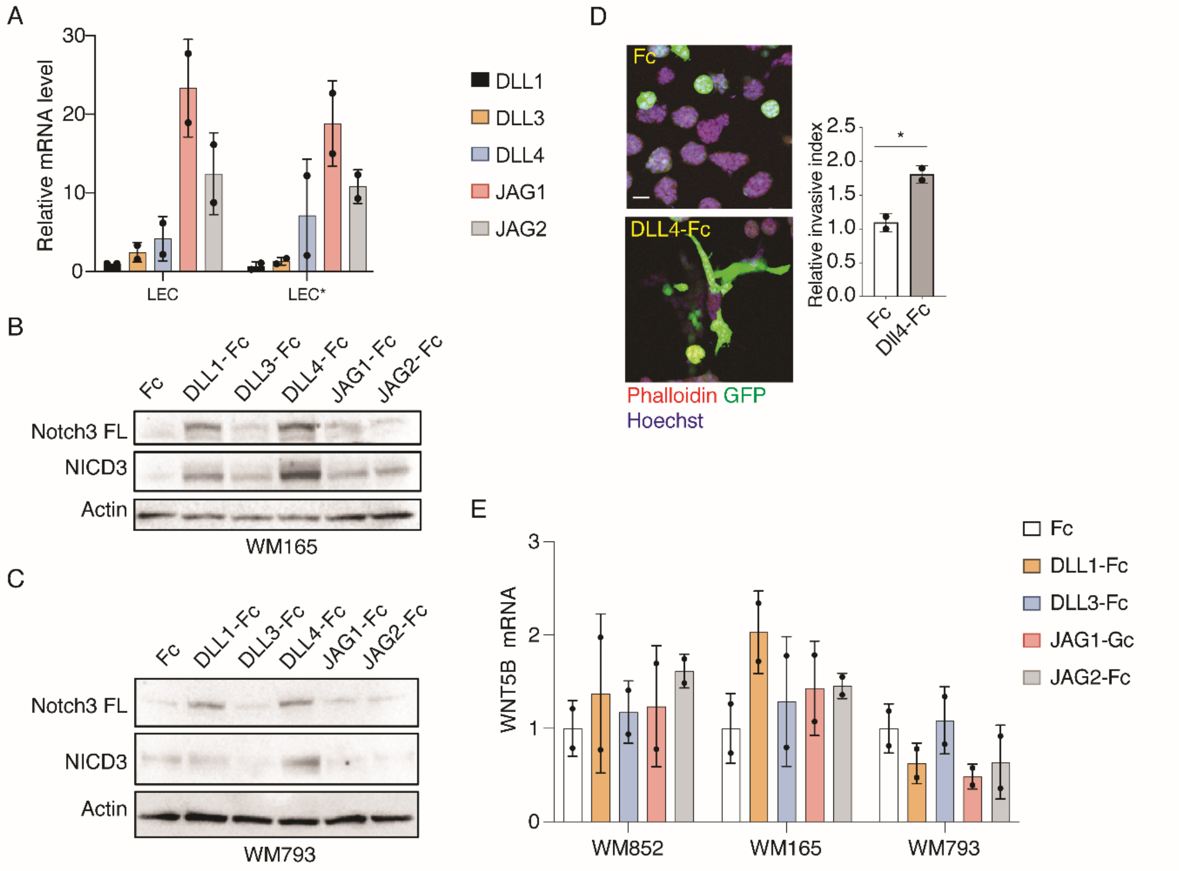


**Supplementary figure 6. DLL4 induces Notch3 and invasive properties of metastatic WM165 melanoma cells.**

**A)** RT- qPCR of the indicated targets in monotypic (LEC) and WM852 melanoma cell co-cultured LECs (LEC*). n=2. Bars, +/- SD. **B-C)** Immunoblotting of the indicated targets in melanoma cells (FL= full length). Cells were cultured on dishes precoated with Fc or Notch ligand-Fc proteins for two days. Representative blots from three experiments are shown. **D)** A 3D fibrin droplet invasion assay of WM165 cells cultured as in B. GFP-expressing melanoma cells were stained with Phalloidin and Hoechst 33342. Graph shows quantification of the relative invasive index from two independent experiments with at least 50 cell clusters quantified/condition. Scale bar = 20µm. Bars, mean +/- SD. **E)** RT-qPCR of *WNT5B* levels in melanoma cells cultured as in A. n=2. Bars, mean +/- SD. *, P<0.05.
